# FlowPot axenic plant growth system for microbiota research

**DOI:** 10.1101/254953

**Authors:** James M. Kremer, Bradley C. Paasch, David Rhodes, Caitlin Thireault, John E. Froehlich, Paul Schulze-Lefert, James M. Tiedje, Sheng Yang He

## Abstract

The presence of resident microbiota on and inside plants is hypothesized to influence many phenotypic attributes of the host. Likewise, host factors and microbe-microbe interactions are believed to influence microbial community assembly. Rigorous testing of these hypotheses necessitates the ability to grow plants in the absence or presence of resident or defined microbiota. To enable such experiments, we developed the scalable and inexpensive FlowPot growth platform. FlowPots have a sterile peat substrate amenable to colonization by microbiota, and the platform supports growth of the model plant *Arabidopsis thaliana* in the absence or presence of soil-derived microbial communities. Mechanically, the FlowPot system is unique in that it allows for total-saturation of the sterile substrate by “flushing” with water and/or nutrient solution via an irrigation port. The irrigation port also facilitates passive drainage of the substrate, preventing root anoxia. Materials to construct an individual FlowPot total ∼$2. A simple experiment with 12 FlowPots requires ∼4.5 h of labor following peat and seed sterilization. Plants are grown on FlowPots within a standard tissue culture microbox after inoculation, thus the Flowpot system is modular and does not require a sterile growth chamber. Here, we provide a detailed assembly and microbiota inoculation protocol for the FlowPot system. Collectively, this standardized suite of tools and colonization protocols empowers the plant microbiome research community to conduct harmonized experiments to elucidate the rules microbial community assembly, the impact of microbiota on host phenotypes, and mechanisms by which host factors influence the structure and function of plant microbiota.

## INTRODUCTION

In an 1885 address to the French Academy of Sciences, Louis Pasteur expressed doubts that axenic (i.e., germ-free) animals were capable of survival^1^. Pasteur went on to credit the work of Duclaux, who reported that peas and beans could not thrive in thoroughly sterilized soil and that plants likely require nutritive assistance from bacteria^2^. Pasteur’s skepticism catalyzed the emergence of a new field: Gnotobiology. Two major motivations for establishing axenic experimental systems that also accommodate gnotobiotic (defined microbiota) or holoxenic (undefined microbiota) hosts are: 1) to elucidate the impact of host-associated microbiota on the host and *vice versa* and 2) to identify host and environmental factors that influence microbiota structure and function. Careful consideration must be taken when designing an axenic growth system to minimize artifacts and sample variability, but not at the expense of versatility.

Environmental factors are the major drivers of differential taxonomic composition and diversity among host-associated microbiota^3–5^. Thus, abiotic factors must be accounted for when performing a microbial colonization experiment for reproducibility of host phenotypes and community dynamics. For example, in the model plant Arabidopsis, humidity influences microbiota composition and the plant’s ability to defend against foliar pathogens^6^. Ecological drivers of soil microbiota composition are associated with physical and chemical factors, including, but not limited to: porosity and water retention, carbon, nitrogen and phosphate availability, pH, percent organic content, and cation exchange capacity^7,8^. When conducting a microbiota colonization experiment and comparing gnotobiotic or holoxenic plants to axenic plants, it is essential to consider abiotic factors of the substrate as well as the growth environment, particularly if the intention of the experiment is to recapitulate a microbial community reminiscent of a soil microbiota.

### Existing growth methodology for plant microbiome research

Axenic Arabidopsis growth can be accomplished with routine tissue culture methodology on a phytonutrient agar substrate (or similar) contained within a light- and gas-permeable vessel. However, routine tissue culture systems do not provide a soil like growth substrate for microbial colonization. Furthermore, agar-based systems are notorious for non-uniform nutrient and O_2_ delivery over time^9^. Hydroponic and aeroponic systems can alleviate issues with nutrient uniformity and O_2_-delivery by agitation and media replenishment, but such systems still do not provide a soil-like growth substrate for microbial colonization^10^. Furthermore, it can be challenging to maintain axenic conditions or prevent cross-contamination in common-reservoir hydroponic systems.

Non-soil substrates, such as sand, quartz, vermiculite and calcined clay, are frequently used in gnotobiotic systems^11–14^. These substrates are porous, thus providing surface area for microbial colonization and root penetration. However, batch-to-batch variation of ceramic substrates can result in a wide range of labile ions^14^. Calcined clay, for example, has sorptive properties that can reduce labile concentrations of P, Fe, Cu and Zn, and desorptive properties that can cause excess labile B, Mg, Ca, S, K, and Mn, leading to potential toxicity of the latter^15^. While thorough washing or soaking of the non-soil substrate can reduce the initial excess of labile ions, flow and drainage are important to reduce significant changes in chemistry over time. Furthermore, the aforementioned non-soil substrates lack significant organic carbon typical of soil, which is typically beneficial to plant and microbial survival and/or growth, unless supplemented.

Soil has been used as a substrate in axenic systems, but often presents challenges, such as contamination or hindered plant growth due to suboptimal sterilization procedures. Numerous sterilization methods have been used with soil, including: autoclaving, dry heat, irradiation, microwave, fumigation by gaseous chemicals, and saturation with various sterilants (i.e., mercuric chloride, sodium azide, formaldehyde)^16^. All methods of soil sterilization alter, to some extent, physical and/or chemical properties of soil. Chemical sterilization methods are not appropriate for plant growth systems due to phytotoxic effects. Autoclaving soil has been shown to increase levels of water-soluble carbon, some ions and reduce pH^17^, but not significantly alter ion exchange capacity. Gamma-irradiation has been reported to minimally disrupt the physical attributes of soils, but can result in the generation of reactive oxygen species, capable of depolymerizing the C-C bond of polysaccharides^18^. Both autoclaving and gamma irradiation can result in changes of the physical structure of the soil, exposing more surface area and thus altering sorptive properties. Complete sterilization of some soils can be achieved with minimal chemical alterations by autoclaving a thin layer of soil for three short (<45 min) autoclave cycles with 18-24 hour intervals^19–21^. Subsequent flushing of sterile substrate can increase plant productivity, presumably by rinsing away soluble phytotoxic byproducts.

### Features and usage of the FlowPot apparatus

Here, we describe an easily-constructed axenic growth system with a peat-based substrate that we call the FlowPot system. We provide a protocol that details how to grow Arabidopsis in axenic FlowPots (no viable microorganisms detected), gnotobiotic FlowPots (inoculated with a defined community of bacteria), and holoxenic FlowPots (inoculated with undefined microbiota extracted directly from natural soil). The peat-based FlowPot system features an inoculation port on each vessel that enables substrate rinsing to remove soluble byproducts of sterilization, provides drainage, and accommodates homogenous inoculation with microbiota and/or nutrients. For added versatility, a mesh retainer allows FlowPots to be inverted for a variety of downstream applications, including dip inoculation or vacuum infiltration of aerial plant tissues, as well as root-specific inoculation. The mesh retainer also prevents the substrate from washing out during the rinsing step. The FlowPot system enables researchers to perform in-depth studies on the phenotype of axenic plants as compared to plants colonized by soil-derived microbiota under highly controlled environmental parameters. The precision, versatility, and robustness of the Flowpot system provides a “germ-free” plant model analogous to the germ-free mouse model. The FlowPot system should be an effective platform to study plant-microbiota interactions *in situ*, including bacterial competitions within a synthetic root microbiota, as recently exemplified by Hassani et al^22^.

### Experimental design

In addition to the microbiota colonization experiment by Hassani et al^22^, here we provide another example experiment that compares the abundance of select photosynthesis-associated proteins in holoxenic and axenic Arabidopsis.

**Assembly protocol**

## MATERIALS

REAGENTS
- Sterile, reverse-osmosis filtered water
- Multi-Terge™ detergent (EMD Millipore, cat. no. 65068); diluted to 2% (v/v)
- Spor-Klenz disinfectant (Steris, USA, cat. no. 652026); dilute to 1% (v/v)
- Linsmaier & Skoog (LS) Medium with Buffer (Caisson Labs, cat. no. LSP03-1 LT);
- Sample containing input microbiota (e.g., from soil)
- R2A agar medium (DIFCO, cat. no. 218263)
- Ethanol, 100% or 95% (v/v) (Fisher Scientific, cat. no. 04-355-451)

EQUIPMENT
- Luer lock PP syringes, 50 ml (Jensen Global, cat. no. JG50CC-LL)
- Female Luer × female Luer adapter, nylon (Cole-Parmer, cat. no. EW-45502-22)
- Mesh fiberglass “Phiferglass”, 18 × 14 standard charcoal mesh (Phifer Incorporated, cat. no. 3003906)
- Soda-glass beads, 3 mm (Sigma-Aldrich, cat. no. Z265926)
- Microboxes (Combiness, USA), model TPD1600 with XXL+ filter
- Filament tape model 893, 18 mm (Scotch Company)
- Redi-Earth plug and seedling mix (Sun Gro Horticulture, Canada). Contains fine Canadian sphagnum peat moss, vermiculite, dolomitic limestone, and a wetting agent

∘ Note this can be substituted with alternative substrates, but plant performance may vary.

- Medium vermiculite, horticultural grade
- Polypropylene trays (United Scientific Supplies, cat. no. 81701)
- Cable ties, 22 mm (TENAX Corporation, Baltimore, USA, cat. no. 120094)
- Sun bag (Sigma-Aldrich, cat. no. B7026)
- Whirl-Pak bags (Nasco, cat. no. 01-812-6C)
- Cell strainer, 40 µm (Falcon, cat. no. 352340)
- Drill bit, 8.8 mm (e.g., Chicago-Latrobe, cat no. 47329)
- Blocks of polypropylene, 12 cm × 8 cm × 1 cm (United States Plastic Corp, cat. no. 42605); alternatively, use Ranin RT-L1000 tip box inserts

## GENERAL EQUIPMENT

- Biosafety cabinet or laminar flow hood (e.g., Logic+ Class II A2 Biological Safety Cabinet, Labconco, cat. no. 302611100; or Console Horizontal Airflow Workstation, Nuaire, cat. no. NU-301-530)
- Standard Sterile Petri plates, 100 × 15 mm (VWR, cat. no. 25384-302)
- Test tube clamp or clamp modified hemostat (e.g., Stoddard Clamp, United Scientific Supplies, cat. no. TTCL03)
- Pipet and 1 ml filter tips (e.g., classic PR-1000 pipette and 1 ml RT-L1000F filter tips, Rainin, cat. nos. 17008653 and 17002920)
- Funnel, 150 mm (Fisher Scientific, cat. no. 10-500-3)
- Glass Erlenmeyer flasks, 2 L (Corning, cat. no. 4980-2L)
- Sterile glass media bottles with screw cap, 2 L (Corning, cat. no. 1395-2L)
- Sterile graduated cylinders, 500ml (Thermo Scientific, cat. no. 36620500)
- Bunsen burner (e.g., Humboldt Manufacturing Company, cat. no. H5870)
- Test tube racks (Thermo Scientific, cat. no. 59700020)
- Miter saw (e.g., Ryobi, cat. no. DC970K-2)
- Drill (e.g., 18-Volt Compact Drill/Driver, Dewalt, cat. no. DC970K-2)
- Growth chamber with desired lighting (e.g., Percival cat. no. CU36L5)

## REAGENT SETUP

- Dilute Multi-Terge detergent (EMD Millipore) to 2% (v/v)
- Dilute Spor-Klenz disinfectant concentrate (Steris, USA) to 1% (v/v)
- **LS** One packet of powder purchased from Caisson Labs, when dissolved in 1 L of deionized water, will result in a 1x concentrate of LS at pH 5.7 at 25 °C. Autoclave the solution. This can be stored at room temperature (22-25 °C) after sterilization for months. A 1x concentrate of LS from Caisson Labs contains: NH_4_NO_3_ 1650 mg/L, H_3_BO_3_ 6.2 mg/L, CaCl_2_ 332.2 mg/L, CoCl_2_. 6H_2_O 0.025 mg/L, CuSO_4_. 5H_2_O 0.025 mg/L, EDTA disodium dihydrate 37.26 mg/L, MES 200 mg/L, MgSO_4_ 180.7 mg/L, MnSO_4_. H_2_O 16.9 mg/L, Na_2_MoO_4_. 2H_2_O 0.25 mg/L, Myo-Inositol 100 mg/L, KHCO_3_ 98 mg/L, KI 0.83 mg/L, KNO_3_ 1900 mg/L, KH_2_PO_4_ 170 mg/L, Thiamine hydrochloride 0.4 mg/L, ZnSO_4_. 7H_2_O 8.6 mg/L, NaCl 0.138 M, KCl 0.0027 M.
- **R2A** Weigh out 18.2 g of powder in 1 L of water. Mix thoroughly. Autoclave at 121°C for 20 minutes on liquid cycle, and pour into sterile Petri dishes. R2A media from Difco contains (in g/L): yeast extract 0.5, proteose peptone No.3 0.5, casamino acids 0.5 g, dextrose 0.5, soluble starch 0.5, sodium pyruvate 0.3 g, dipotassium phosphate 0.3, magnesium sulfate 0.05, agar 15.0.

## PROCEDURE

### Construction of FlowPots • TIMING ∼1.5 h

Note: FlowPots are reusable. This phase of the protocol only needs to be performed for initial construction.

1. Remove the pistons from the 50 ml polypropylene (PP) Luer taper syringes. Using a miter saw with a fine-tooth blade, cut the syringes at the “20 ml” mark, retaining only the portion with the Luer connector. Mount the blade on the miter saw backwards for a smoother cut, and sand if needed. Remove any residual shards with a vacuum and a moist cloth. Soak the syringe tops for 20 minutes in 2% Multi-Terge ionic detergent, and subsequently rinse the syringe tops in distilled water to remove all traces of the detergent. Autoclave prior to FlowPot construction.

a. Note: avoid syringes that have have silicon oil or other lubricants within the barrel.
b. Caution: use proper safety precautions when using the miter saw, and proper eye protection when cutting plastic.
2. Cut 5 × 5 cm squares of mesh fiberglass. Autoclave prior to FlowPot construction.
3. Rinse 3 mm soda-glass beads 6 times with distilled water. Dry and autoclave prior to FlowPot construction.
4. To construct a FlowPot stand, drill four holes in a 12 × 8 × 1 cm block of autoclave-compatible plastic (polypropylene or polycarbonate) using an 8.8 mm drill bit. Orient the holes so they are evenly distributed with adequate spacing from stand edge so that the FlowPots do not exceed the stand boundaries.

a. Caution: Use proper safety precautions when drilling inserts and using power tools.
b. Note: We routinely use disposable the inserts from Ranin RT-L1000 pipette tip boxes for this purpose.
c. Note: Although a stand of these dimensions can accommodate seven FlowPots, in our experience, plant growth is optimal with four FlowPots.

### Preparation of the soil extract source microbiota • TIMING varies

Note: procurement of a source microbiota can be performed in advance of the experiment and modified depending on the input community characteristics and experimental parameters of your choosing. As an alternative to a soil source community, one can use defined communities of microorganisms derived from pure cultures^22^.

5) For soil source communities, remove topsoil and collect soil >6 cm below the surface. Let soil sit for 1 week at room temperature with ∼50% relative humidity, and sift through a 3 mm^2^ galvanized steel screen to remove large debris. Aliquot soil in 100 g increments and store at 4°C in Whirl-Pak bags.

### Sterilization of the substrate • TIMING ∼3 d

6) Blend a 1:1 ratio of peat potting mix and medium vermiculite (substrate). Moisten with distilled water to achieve moisture content of approximately 60%. Evenly distribute the substrate on clean polypropylene laboratory trays at a depth of approximately 2 cm. Cover the surface of each tray with aluminum foil or autoclave paper in such a way that liquid will not collect on top during autoclaving and flow onto the substrate. Autoclave for 30 minutes on liquid cycle (121.1 °C, 15 PSI, slow exhaust with forced liquid cooling) and bring to room temperature immediately after autoclaving.

a. Note: Do not let materials sit in the autoclave after cycling because this may cause the substrate to dry out, resulting in increased hydrophobicity and suboptimal plant growth.
7) Homogenize substrate in a sterile container and subsequently distribute on polypropylene laboratory trays. Let sit covered with aluminum foil or autoclave paper at room temperature (22-25 °C) for 24-48 hours.
8) Autoclave the substrate a second time (to kill any spores) for 30 minutes on liquid cycle (121.1 °C, 15 PSI, slow exhaust with forced liquid cooling). Pre-clean the surface of a laminar flow hood using Spor-Klenz. Immediately after autoclaving, place the autoclaved trays of substrate in the pre-cleaned laminar flow hood and bring to room temperature. Once at room temperature, aseptically homogenize the substrate in the sterile laminar flow hood. Cover the trays of substrate with sterile aluminum foil or autoclave paper. Leave covered at room temperature (22-25 °C) for 24-48 hours.

a. Note: Depending on the moisture content of your substrate, relative humidity, and the calibration of your autoclave, autoclave parameters may need to be optimized to ensure sterility whilst preserving the integrity of the substrate.
b. Caution: Spor-Klenz is caustic and an eye/skin irritant. Use proper safety precautions.

### Seed preparation • TIMING ∼3 d

9) Sterilize seeds using the vapor-phase sterilization protocol (chlorine gas) for 6-8 hrs^23^. Check for seed-borne contaminants and germination efficiency by incubating an aliquot of seeds on R2A agar for 1 week at 22°C in the dark.

a. Caution: chlorine gas is caustic! Use proper safety precautions.
b. Caution: chlorine gas will dissolve many types of permanent ink. Be sure to label samples with a solvent resistant marker or use laminated labels.
c. Note: It is important to open sterile seed aliquots in a sterile laminar flow hood for at least 10 minutes after sterilization to adequately remove residual chlorine gas.
d. Note: aliquots of seed can be sterilized in bulk and stored under appropriate conditions for future use.
10) Allow seeds to imbibe during a 48-hour stratification period in sterile, distilled water at 4°C in the dark prior to sowing.

a. Note: check for effective decontamination of microorganisms by spreading an aliquot of seeds on R2A media and monitor for microbial growth after one week.

### Assembly of FlowPots • TIMING ∼2 h

11) Aseptically place 10 sterile glass beads into each of autoclaved syringe tops. Gently fill each syringe top with the twice-autoclaved substrate mixture until slightly heaping (∼ 0.5 cm). Cover barrel end of the syringe top with the square mesh and secure with a cable tie. Trim the excess edges of the square mesh. Fasten a female-female Luer connector and place on a test tube rack to complete FlowPot construction.

a. Critical: Do not overpack the substrate. Compaction can lead to suboptimal plant growth. Within an experiment, it is critical to maintain the same relative compaction for all FlowPots.
b. Note: To stabilize FlowPots during assembly, while filling, we often use a sterile test tube rack.
c. Note: We recommend using a cable tie gun: Thomas & Betts Ty-Rap Tool (http://www.cableorganizer.com/thomas-betts/ty-rap-tool.html).
12) Once the test tube rack is full, place the test tube rack full of assembled FlowPots in a Sun bag and loosely close the end with autoclave tape such that the risk of contamination is minimized when the bag is removed from the autoclave, yet steam may still permeate the bag during sterilization. Autoclave for 30 minutes on liquid cycle (121.1 °C, 15 PSI, slow exhaust with forced liquid cooling). Immediately after autoclaving, seal the opening of the Sun bag and move to a sterile hood.

a. Note: Alternative autoclave-safe bags can be used instead of Sun bags, and FlowPots can also be autoclaved directly in microboxes as well.
13) Center and fasten the drilled FlowPot stand to the inside bottom a microbox tissue culture vessel (model TPD1600 with XXL+ filter) using filament tape. Autoclave constructed boxes and lids according to the manufacturer’s instructions prior to use.

a. Note: We routinely use 18 mm filament tape model 893 (Scotch, USA), but alternative tapes are suitable.

**? TROUBLESHOOTING**

### FlowPot irrigation and inoculation • TIMING ∼2 h

14) Add 950 ml of sterile distilled H_2_O and 50 g of sieved soil to a sterile 2-L Erlenmeyer flask. Agitate soil slurry on a rotary shaker for 20 minutes at 22°C at ∼200 rpm, and subsequently let settle for 5 minutes. Filter the supernatant through a 40 µm cell strainer into a sterile 2-L Nalgene media bottle.

a. Note: Allowing the slurry to settle increases reproducibility of colonization^24^ and reduces filter clogging.
15) Divide the soil slurry into two. Prepare the holoxenic inoculum by directly mixing the strained soil slurry with equal parts 1x LS media. To prepare a sterile mock inoculum, autoclave the remaining portion of the strained soil slurry for 45 minutes (121.1 °C, 15 PSI, slow exhaust with forced liquid cooling), then mix with equal parts 1x LS media in a sterile laminar flow hood, bringing the final concentration of LS to 1/2x.

a. Note: The amount of inoculum needed for each condition will be determined by the number of FlowPots being prepared.
16) Using a sterile test tube clamp, grasp each FlowPot and invert over a sterile funnel placed atop a waste flask. While inverted, use a sterile 50 ml syringe to slowly, and aseptically infiltrate each FlowPot via the Luer connector with 50 ml of sterile H_2_O. Apply even pressure for 30 seconds during the infiltration. After water infiltration, place the FlowPot on a sterile test tube rack. To reduce the risk of contamination, we recommend ethanol-flaming the test tube clamps between each FlowPot infiltration. The preparation of axenic FlowPots should be performed separately from those that are holoxenic and in a Biosafety cabinet or laminar flow hood.

a. Caution: Use proper safety precautions when working with fire and flammable solvents such as ethanol.
b. Note: Occasionally, the glass beads are oriented in a way that the infiltration port is obscured. In this case, a sterile syringe needle may be inserted into the infiltration port to clear the blockage.
c. Note: An alternative to the test tube clamp holder is a modified hemostat with semicircular stainless steel bands to grip the FlowPots.

**? TROUBLESHOOTING**

17) Let water-infiltrated FlowPots sit for 30 minutes, then infiltrate the FlowPots with the desired input community mixture. Evenly mix the input community prior to infiltration.
18) Place irrigation port of inoculated FlowPots in the drilled holes of the FlowPots stand within the sterile microboxes. We recommend 4 FlowPots per microbox.

### Sowing seeds • TIMING ∼0.5 h

19) Aseptically sow approximately 8 seeds per FlowPot using a pipette with filter tips.
20) Place microboxes with planted FlowPots in the appropriate growth conditions.

a. Note: After sowing, make sure the microbox lids are completely sealed to maintain consistent humidity and sterility.
b. Note: For *Arabidopsis thaliana* Col-0, we recommend the following growth conditions: 23 °C with 12/12 day/night light cycles at of ∼80 µE m^−2^ s^−1^.
21) Aseptically thin boxes to 3 plants per pot using flamed forceps 7-10 days after germination.

a. Critical step: Check sterility by placing thinned plants on R2A agar and incubate for at least 7 days at 22 °C.

**? TROUBLESHOOTING**

### Example experiment: quantification of photosynthesis-associated proteins

#### • TIMING ∼5 h

For this experiment, Michigan Soil (named “MS13MSU”) was collected during Fall 2013 from an sandy loam agricultural field at Michigan State University, East Lansing, Michigan, USA 42.709°N, −84.466°E). The field was used for *Miscanthus* cultivation, and had not been subjected to tilling, fertilization, or any intervention for a minimum of 10 years. Soil was collected from 5 to 15 cm below the surface. Upon collection, soil was spread out on tables and allowed to sit for 1 w at room temperature with ∼50% relative humidity. Large debris was removed using a 3 mm galvanized metal screen, and 50 g aliquots were prepared in Whirl-Pak bags and stored at 4°C in the dark.

For leaf protein extraction single Arabidopsis leaves (3 week old) from FlowPot-grown axenic or holoxenic plants (MS13MSU community) were collected and homogenized using Minute-Chloroplast Isolation Kit (Invent Biotechnology, Inc. Plymouth, MN, USA) according to the manufacturer’s protocol. Chlorophyll content of total leaf homogenate was determined by the method of Arnon^25^ to give a final concentration of 1 mg chlorophyll/ml for all samples. All total leaf homogenate (5 mg chlorophyll total) samples were solubilized in Laemmli buffer ^26^ and resolved by SDS-PAGE and either stained with Coomassie Blue or transferred onto polyvinylidene difluoride (PVDF) membrane (Invitrogen) and probed with the following antibodies: anti-D1 Protein (Agrisera, AS01 016), OEC33 (Agrisera, AS06 142-33), OEC23 (Agrisera, AS06 167) or antibodies produced in house: Anti-Toc159, Anti-Toc75, anti-Tic110, anti-ClpC^27^. All primary antibodies were incubated using a 1:4,000 dilution. The detection method employed used a secondary anti-rabbit conjugated to alkaline phosphatase (KLP, Inc. Gaithersburg, MD) at 1:5,000 dilution for 1 hour in 5% DM/TBST. The blots were developed using a standard Alkaline Phosphatase (AP) detection system with BCIP/NBT as substrates (Sigma-Aldrich, St. Louis, MO, USA). We found that holoxenic and axenic Arabidopsis had similar levels of examined photosynthesis-associated proteins (Fig. 2), which is in line with the healthy appearance of axenic Arabidopsis.

**Figure 1:**
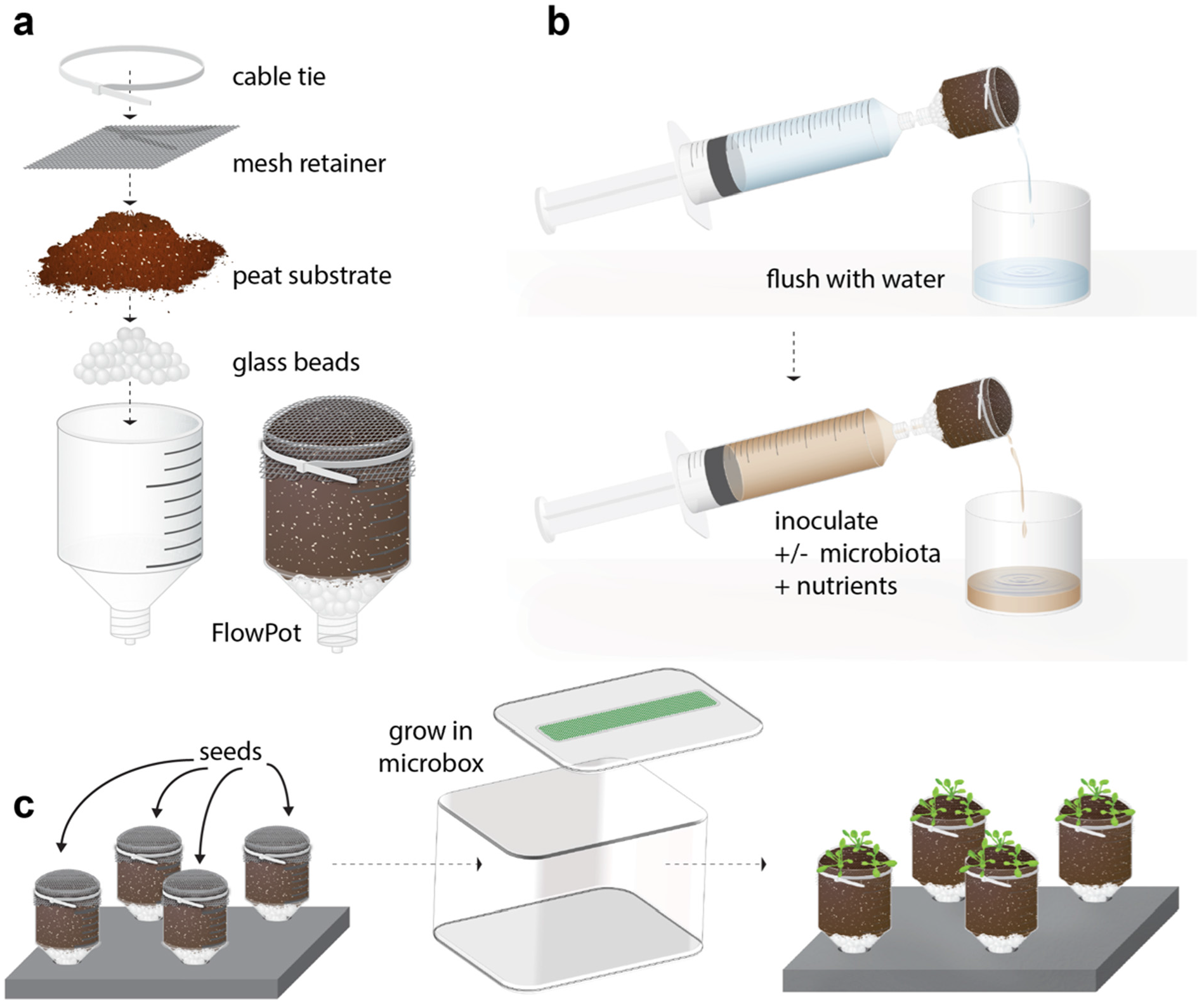
Diagram of FlowPot Assembly. Each FlowPot is prepared by (**a**) adding glass beads to the Luer end of a truncated syringe, followed by the addition of twice-autoclaved peat, covered with a mesh retainer, and then secured with a cable tie. Assembled FlowPots are then autoclaved, (**b**) aseptically irrigated with sterile water, and inoculated with nutrients and any desired input microbiota. (**c**) Decontaminated Arabidopsis seeds, stratified and imbibed with water, are sown onto each FlowPot. FlowPots are then placed into microboxes on stands Stratified. Imbibed sterile Arabidopsis seed is sown onto each FlowPot and FlowPots are placed into sterile microboxes. The microboxes containing FlowPots are placed in a growth chamber or greenhouse with desired lighting and temperature conditions for plant growth.

**Figure 2.**
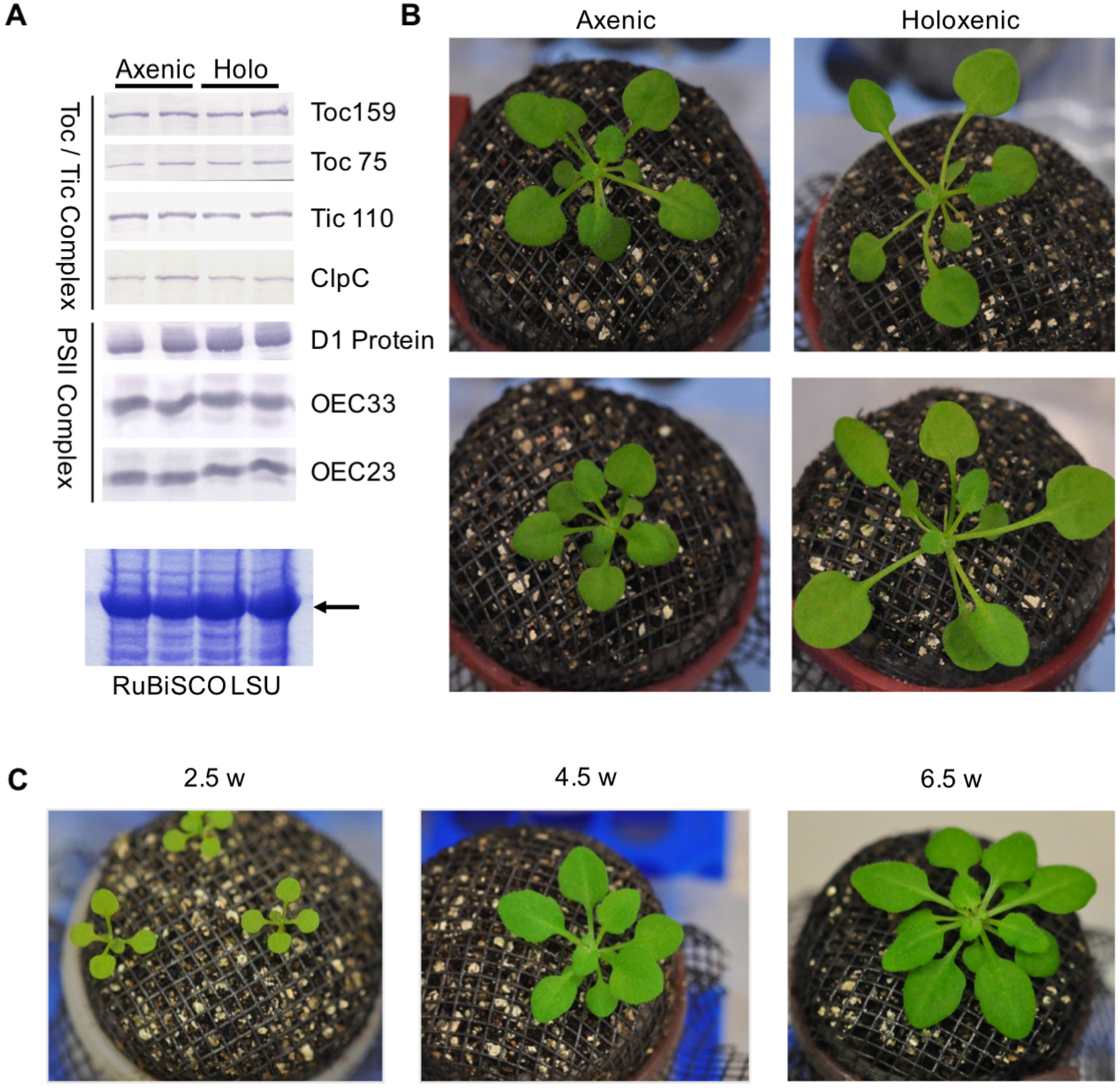
*Arabidopsis thaliana* grown in FlowPots with axenic or holoxenic substrate, and photosynthetic protein detection from leaf tissue. (A) Photosynthesis-associated protein quantification from total protein extracts of rosette tissue 3 weeks post germination. (B) Arabidopsis in FlowPots 4 weeks post germination. Holoxenic substrate was inoculated with the “MS13MSU” community. (C) Axenic Arabidopsis growth, photographed at 2.5 weeks, 4.5 weeks and 6.5 weeks post germination. Rosette images and the protein gel are representative of at least three replicated experiments.

**? Troubleshooting**

Troubleshooting advice can be found in **Table 1.**

**Table 1.**
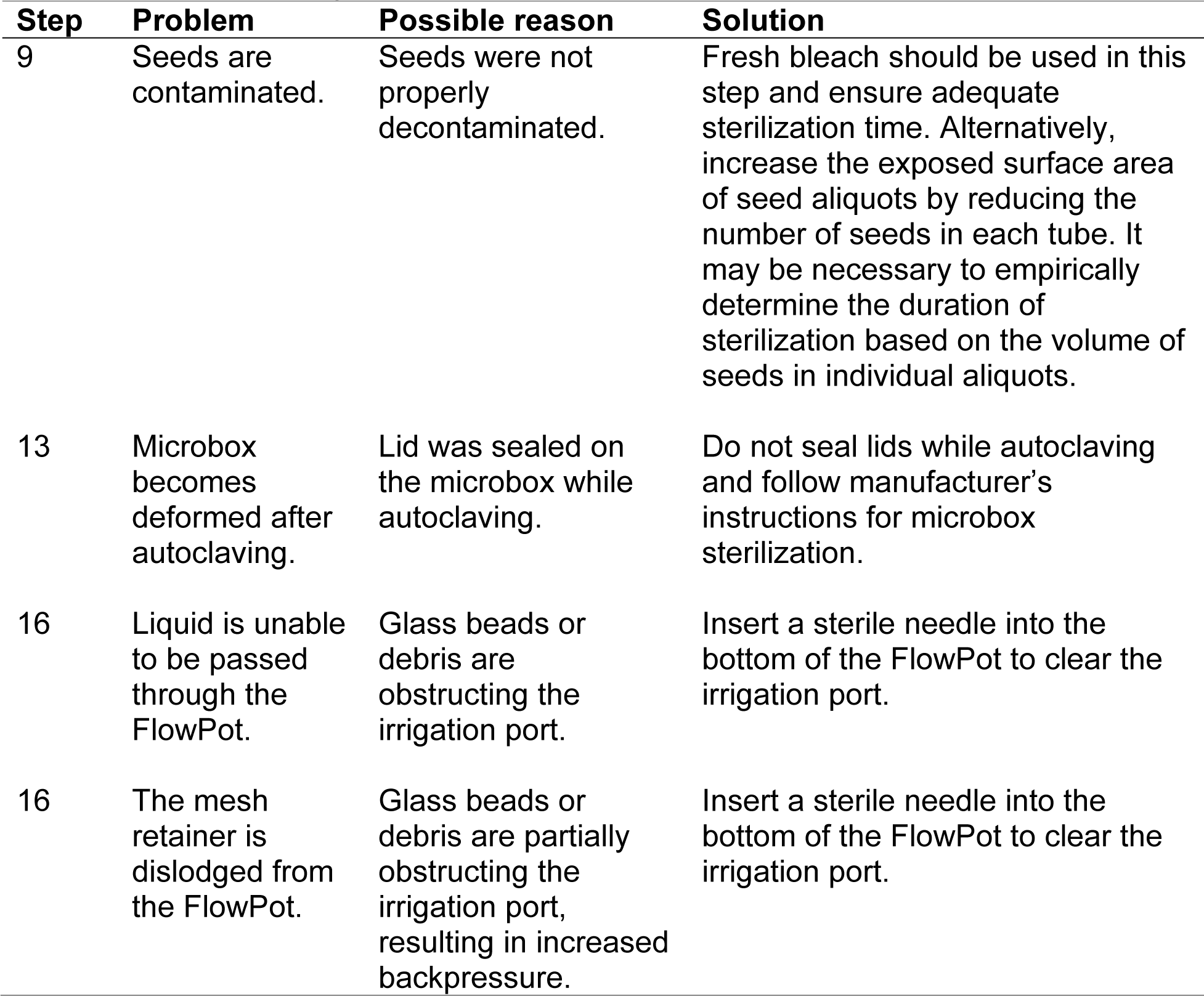
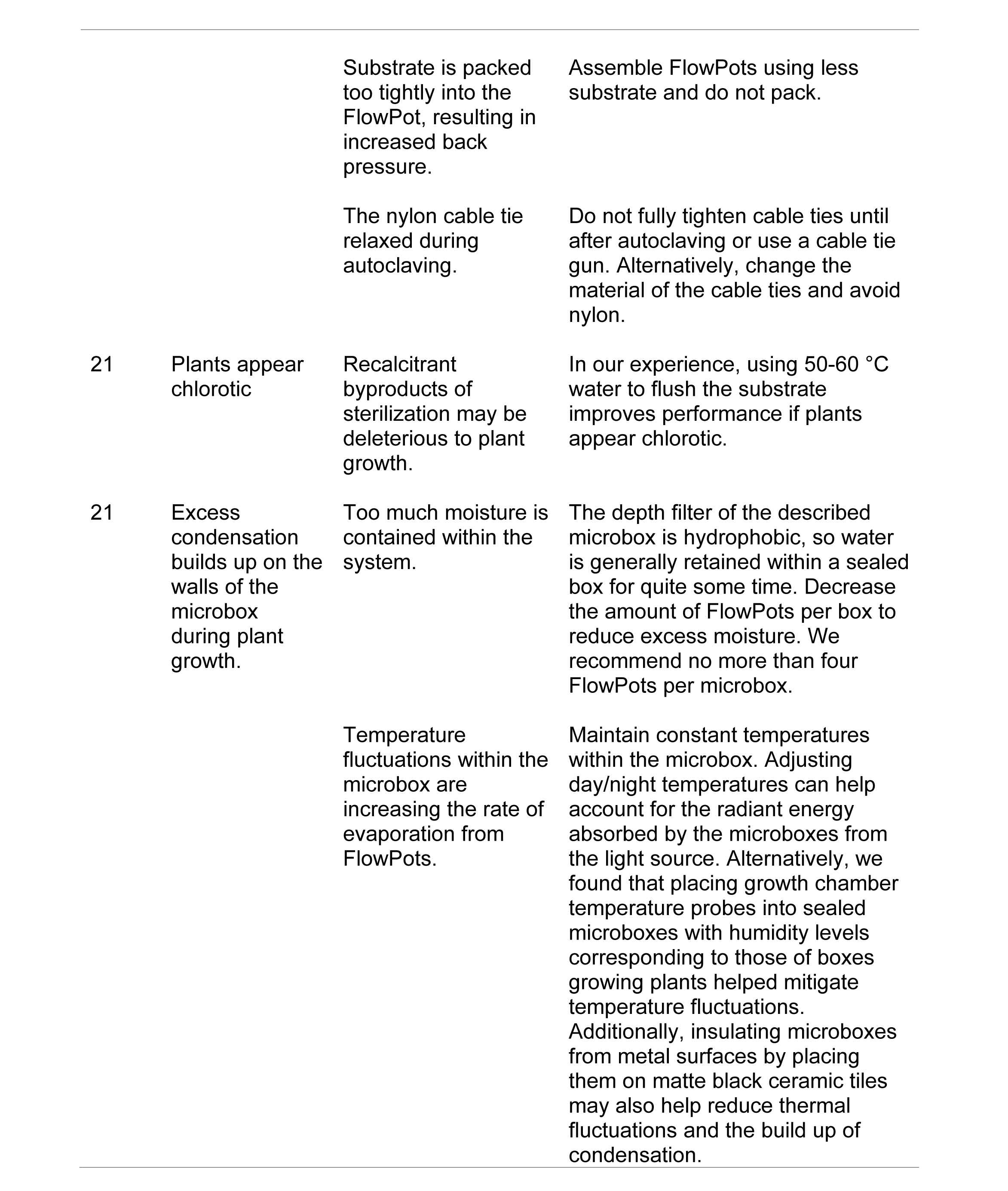
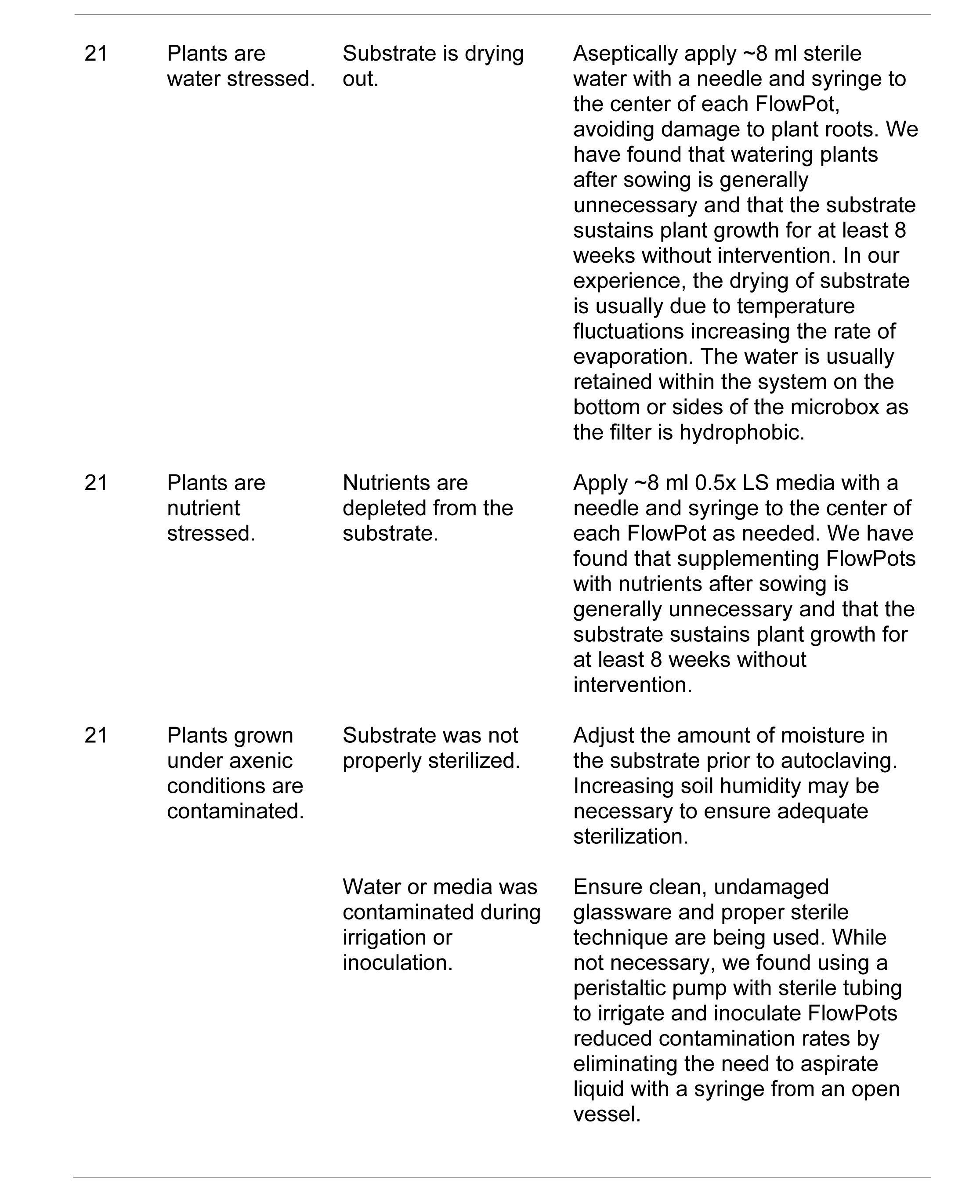
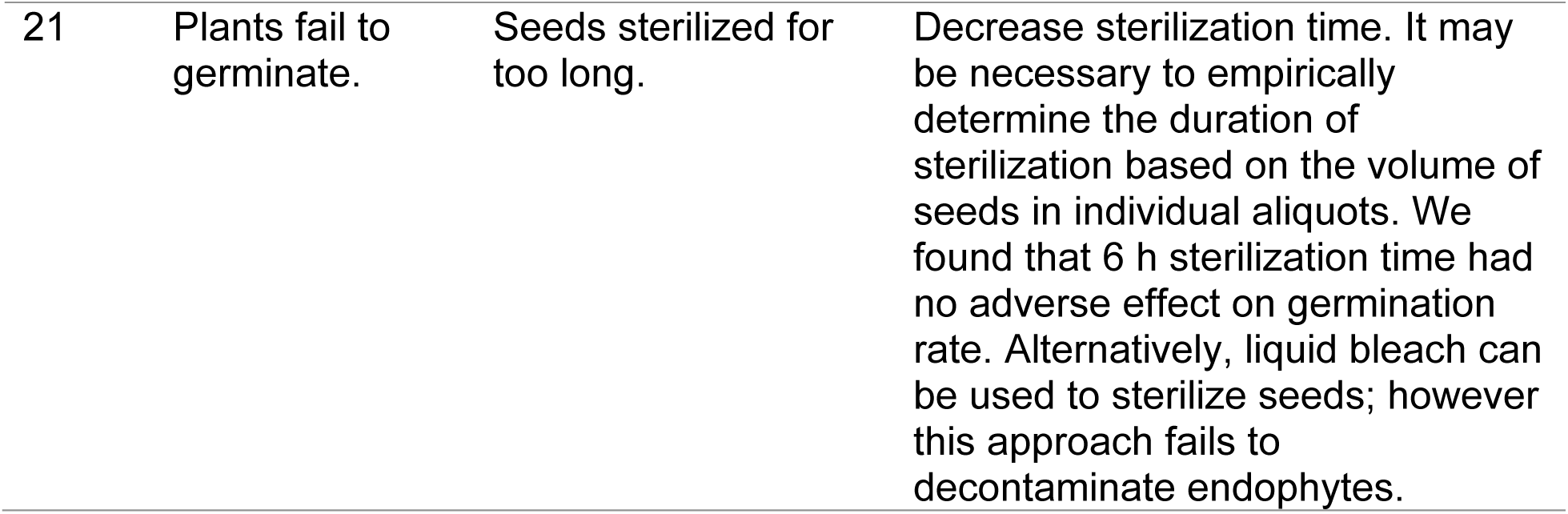
Troubleshooting table.

| Step | Problem | Possible reason | Solution |
| --- | --- | --- | --- |
| 9 | Seeds are contaminated. | Seeds were not properly decontaminated. | Fresh bleach should be used in this step and ensure adequate sterilization time. Alternatively, increase the exposed surface area of seed aliquots by reducing the number of seeds in each tube. It may be necessary to empirically determine the duration of sterilization based on the volume of seeds in individual aliquots. |
| 13 | Microbox becomes deformed after autoclaving. | Lid was sealed on the microbox while autoclaving. | Do not seal lids while autoclaving and follow manufacturer's instructions for microbox sterilization. |
| 16 | Liquid is unable to be passed through the FlowPot. | Glass beads or debris are obstructing the irrigation port. | Insert a sterile needle into the bottom of the FlowPot to clear the irrigation port. |
| 16 | The mesh retainer is dislodged from the FlowPot. | Glass beads or debris are partially obstructing the irrigation port, resulting in increased backpressure. | Insert a sterile needle into the bottom of the FlowPot to clear the irrigation port. |
|  |  | Substrate is packed too tightly into the FlowPot, resulting in increased back pressure. | Assemble FlowPots using less substrate and do not pack. |
|  |  | The nylon cable tie relaxed during autoclaving. | Do not fully tighten cable ties until after autoclaving or use a cable tie gun. Alternatively, change the material of the cable ties and avoid nylon. |
| 21 | Plants appear chlorotic | Recalcitrant byproducts of sterilization may be deleterious to plant growth. | In our experience, using 50-60 °C water to flush the substrate improves performance if plants appear chlorotic. |
| 21 | Excess condensation builds up on the walls of the microbox during plant growth. | Too much moisture is contained within the system. | The depth filter of the described microbox is hydrophobic, so water is generally retained within a sealed box for quite some time. Decrease the amount of FlowPots per box to reduce excess moisture. We recommend no more than four FlowPots per microbox. |
|  |  | Temperature fluctuations within the microbox are increasing the rate of evaporation from FlowPots. | Maintain constant temperatures within the microbox. Adjusting day/night temperatures can help account for the radiant energy absorbed by the microboxes from the light source. Alternatively, we found that placing growth chamber temperature probes into sealed microboxes with humidity levels corresponding to those of boxes growing plants helped mitigate temperature fluctuations. Additionally, insulating microboxes from metal surfaces by placing them on matte black ceramic tiles may also help reduce thermal fluctuations and the build up of condensation. |
| 21 | Plants are water stressed. | Substrate is drying out. | Aseptically apply ~8 ml sterile water with a needle and syringe to the center of each FlowPot, avoiding damage to plant roots. We have found that watering plants after sowing is generally unnecessary and that the substrate sustains plant growth for at least 8 weeks without intervention. In our experience, the drying of substrate is usually due to temperature fluctuations increasing the rate of evaporation. The water is usually retained within the system on the bottom or sides of the microbox as the filter is hydrophobic. |
| 21 | Plants are nutrient stressed. | Nutrients are depleted from the substrate. | Apply ~8 ml 0.5x LS media with a needle and syringe to the center of each FlowPot as needed. We have found that supplementing FlowPots with nutrients after sowing is generally unnecessary and that the substrate sustains plant growth for at least 8 weeks without intervention. |
| 21 | Plants grown under axenic conditions are contaminated. | Substrate was not properly sterilized. | Adjust the amount of moisture in the substrate prior to autoclaving. Increasing soil humidity may be necessary to ensure adequate sterilization. |
|  |  | Water or media was contaminated during irrigation or inoculation. | Ensure clean, undamaged glassware and proper sterile technique are being used. While not necessary, we found using a peristaltic pump with sterile tubing to irrigate and inoculate FlowPots reduced contamination rates by eliminating the need to aspirate liquid with a syringe from an open vessel. |
| 21 | Plants fail to germinate. | Seeds sterilized for too long. | Decrease sterilization time. It may be necessary to empirically determine the duration of sterilization based on the volume of seeds in individual aliquots. We found that 6 h sterilization time had no adverse effect on germination rate. Alternatively, liquid bleach can be used to sterilize seeds; however this approach fails to decontaminate endophytes. |

#### • TIMING

**Pre-experiment preparation**:

Steps 1-4, construction of FlowPots: ∼1.5 h

Step 5, preparation of the soil extract source community: varies

**Days 1-4**:

Steps 6-8, sterilization of the substrate: ∼3 d

Steps 9-10, seed preparation: ∼3 d

**Day 4**:

Steps 11-13, assembly of FlowPots: ∼2 h

Steps 14-19, FlowPot irrigation and inoculation: ∼2 h

Steps 20-21, sowing seeds: ∼0.5 h

### Anticipated Results

We have shown that the FlowPot system can be used to study microbiota dynamics *in planta*^22^ and for basic characterization of plant phenotypes (Fig 2). In the future, this system should allow researchers to investigate a wide variety of other questions in plant and microbial sciences where presence and absence of individual microbes or synthetic or natural microbiota are a critical part of experimentation. These could include study of the rules governing microbial community assembly, the impact of microbiota on host phenotypes, gene expression and epigenetic processes, and mechanisms by which host factors influence the structure and function of plant microbiota. Specific experimental designs to address such questions may necessitate modification of certain aspects of the FlowPot system, which will likely lead to specialized iterations of the basic system described here.

## Acknowledgements

We would like to thank Caleigh Griffin, Alec Bonifer, Alan Mundakkal, and Franchesca Dion for assistance with FlowPot assembly and workflow optimization, Dr. M. Amine Hassani and Dr. Stephane Hacquard for critical reading and helpful comments on this manuscript, and Dr. Brian Kvitko and Dr. JP Jerome for their contributions to growth system development. This project was supported by funding from Gordon and Betty Moore Foundation (GBMF3037), and the Department of Energy (the Chemical Sciences, Geosciences, and Biosciences Division, Office of Basic Energy Sciences, Office of Science; DE-FG02-91ER20021 for infrastructural support).

## Author Contributions

J.M.K, JT and S.Y.H designed the experiments. J.M.K, B.P, D.R., C.T. and J.F. performed and/or analyzed the experiments. P.S.-L supervised J.M.K. during protocol optimization at Max Planck Institute for Plant Breeding Research, Cologne. J.M.K and S.Y.H wrote and finalized the manuscript with input from all co-authors.

